# Quantifying defensive behavior and threat response through integrated headstage accelerometry

**DOI:** 10.1101/2021.12.21.473691

**Authors:** Rebecca Younk, Alik S Widge

## Abstract

**Background:** Defensive and threat-related behaviors are common targets of investigation, because they model aspects of human mental illness. These behaviors are typically quantified by video recording and post hoc analysis. Those quantifications can be laborious and/or computationally intensive.

Depending on the analysis method, the resulting measurements can be noisy or inaccurate. Other defensive behaviors, such as suppression of operant reward seeking, require extensive animal pre-training.

**New Method:** We demonstrate a method for quantifying defensive behavior (immobility or freezing) by 3-axis accelerometry integrated with an electrophysiology headstage. We tested multiple pre-processing and smoothing methods, and correlated them against two common methods for quantification: freezing as derived from standard video analysis, and suppression of operantly shaped bar pressing. We assessed these three methods’ ability to track defensive behavior during a standard threat conditioning and extinction paradigm.

**Results:** The best approach to tracking defensive behavior from accelerometry was Gaussian filter smoothing of the first derivative (change score or jerk). Behavior scores from this method reproduced canonical conditioning and extinction curves at the group level. At the individual level, timepoint-to-timepoint correlations between accelerometry, video, and bar press metrics were statistically significant but modest (largest r=0.53, between accelerometry and bar press).

**Comparison with existing methods:** The integration with standard electrophysiology systems and relatively lightweight signal processing may make accelerometry particularly well suited to detect behavior in resource-constrained or real-time applications. At the same time, there were modest cross-correlations between all three methods for quantifying defensive behavior.

**Conclusions:** Accelerometry analysis allows researchers already using electrophysiology to assess defensive behaviors without the need for additional behavioral measures or video. The similarities in behavioral tracking and modest correlations between each metric suggest that each measures a distinct aspect of defensive behavior. Accelerometry is a viable alternative to current defensive measurements, and its non-overlap with other metrics may allow a more sophisticated dissection of threat responses in future experiments.

**Highlights:**

- A novel method to assess defensive behavior and immobility based on headstage accelerometry
- Compatible with readily available, open-source neurophysiology systems
- Provides behavioral insights without the need for video analysis software and with relatively minimal processing, suitable for rapid closed-loop experiments

## Introduction

Defensive behaviors in animals, such as freezing and rapid escape, are commonly used as models of human negative-valence processes (Campos et al., 2013; Colom-Lapetina et al., 2019; Deslauriers et al., 2018). Over-emission or contextually inappropriate emission of these defensive behaviors can model aspects of human psychiatric disorders (Robinson et al., 2019; Terburg et al., 2018). The standard approach to quantifying these behaviors in experimental preparations is through video analysis. Traditionally, these videos are scored using manual observation or automatic classifiers which assess behavior per video frame (Anagnostaras et al., 2010; Sousa et al., 2006). Automatic classifiers are becoming substantially more advanced, incorporating deep learning algorithms to more accurately assess, and in some cases, predict animal behavior (Liu et al., 2021; Mathis et al., 2020; von Ziegler et al., 2021). Although video is the traditional method of behavior analysis, it might not be suitable for all situations. Manual scoring has inter-rater variability and is highly laborious. Results from automatic classifiers can depend on camera position, video quality, frame rate, lighting, and the presence of other wires and cables within the video frame, each of which may distort behavioral results. Traditional video analysis primarily assesses the change in pixels between frames. It does not perform well if there are multiple animals present (e.g., in assessing defensive responses to a conspecific or in other social assays). More advanced classifiers are more robust to these problems, but at the cost of greatly increased computational burden that may make them inaccessible to many labs.

Another common approach to quantifying defensive behavior is bar press suppression (Amorapanth et al., 1999; Li et al., 2014; Mueller et al., 2009). Animals can be shaped on variable-interval schedules until they reliably press a bar at a near-constant rate to receive reward pellets. Acute threat suppresses this operant response, and the duration/amount of interruption is a metric of defensive behavior. Bar press suppression is easy to quantify without any manual observation or subjective interpretation, but can nevertheless be expensive and labor-intensive. Pre-training animals to a reliable press rate can take weeks, and often needs to be repeated after surgery or other experiment-related interventions.

Continuous motion detection offers a solution to these issues, providing real-time analysis capabilities that are not impeded by video related factors. As neural recording electronics have become more miniaturized, it has become possible to incorporate accelerometers (Venkatraman et al., 2007; Zamora et al., n.d.).

For rodent behavioral studies that include electrophysiology, continuous motion sensors offer another analysis approach, but there is no standardized approach to detecting defensive behavior from accelerometry. Here, we demonstrate one, designed for compatibility with an open-source electrophysiology system. We show that video-derived freezing, suppression of operant bar pressing, and accelerometry capture different but related aspects of defensive behaviors. Thus, accelerometry may be a helpful and easily measured tool to complement existing defensive assays.

## Methods

### Animals and Behavior Paradigm

#### Animals

16 adult male and 12 adult female Long Evans Rats weighing 250-350 grams (Charles River, Madison, WI) were used as subjects. Sample size was determined by available animals used for another, ongoing, study, which was powered to detect a difference in behavior between different brain stimulation protocols. Those protocols had no statistically significant effect and thus all rats were combined for this behavior analysis. Rats were initially housed in pairs in plastic home cages for a minimum of 7 days, to allow for acclimation to the facility. Standard rat chow and water were available ad libitum during facility habituation. All experiments occurred during the light cycle with a 14 h on/10 h off cycle (lights on at 600). Following the acclimation period, animals were handled daily for 5 days to minimize handling stress. Rats were then housed individually in plastic home cages. Food was limited to 10 grams per day until the rat was between 85-95% of its starting weight, then fed 15-20 grams per day to maintain weight within this range for the remainder of behavioral procedures. During the first three days of food restriction, sucrose pellets were placed in the home cage to habituate rats to the reward and facilitate bar press learning. All experimental details were approved by the Institutional Animal Care and Use Committee at the University of Minnesota and performed in compliance with the Guide for the Care and Use of Animals. Research facilities were accredited by the American Association for the Accreditation of Laboratory Animal Care.

#### Apparatus

All behavioral training and experiments were conducted in Coulbourn (Allentown, PA) conditioning chambers (30.5 × 24.1 × 21 cm). The conditioning chamber was housed in a sound attenuating box that contained a ventilation fan and was illuminated by a house light and red puck light (Commercial Electric) for recording purposes. The grid floor (1.6 cm spacing, 4.8 mm diameter rods) was used to administer foot shocks. A retractable bar and food trough were located on one of the aluminum walls and the speaker was positioned on the opposite wall. The camera and attached wide angle lens used for recording (Logitech HD Pro Webcam C910; Neewer Digital high definition .45x super wide-angle lens) was positioned outside of the conditioning chamber above the speaker unit looking down through the plexiglass top. Foot shocks were generated by a Lafayette Instruments scrambled grid current generator (Lafayette, IN).

#### Behavior

Rats were trained to bar press for sucrose pellets on an increasing variable interval (VI) schedule (0, 15, 30, 45, 60) for 40 minutes per day. Once they reached a minimum rate of 10 presses per minute on VI-60, they moved on to electrode implantation and a behavioral paradigm post-surgery. Microelectrode bundles as described in Lo et al., (2020) were implanted in the prefrontal cortex and amygdala, and the results of electrophysiologic recordings will be described in a separate paper.

To evoke freezing behavior, rats underwent a tone-shock conditioning protocol (Figure 1B). The experiment consisted of three phases: habituation/conditioning, extinction, and extinction recall. Each day, rats were exposed to a conditioned stimulus (CS). The intertrial interval between each CS varied (65 - 295s), averaging 3 minutes, but was consistent between rats. Electrophysiology and video were recorded throughout each experimental session. The bar for sucrose pellets was available any time the rat was in the chamber and continued to reinforce at VI-60. On day 1, for habituation, rats were presented with five trials of the CS (30s, 82 dB tone). They were subsequently conditioned through seven presentations of the CS + unconditioned stimulus (US; 0.6 mA, 0.5s foot shock) immediately after the CS. Extinction consisted of 20 presentations of the CS in absence of the US, administered in the same chamber as conditioning. On day 3 the CS was presented 6 times without the US to measure extinction memory (recall).

**Figure 1:**
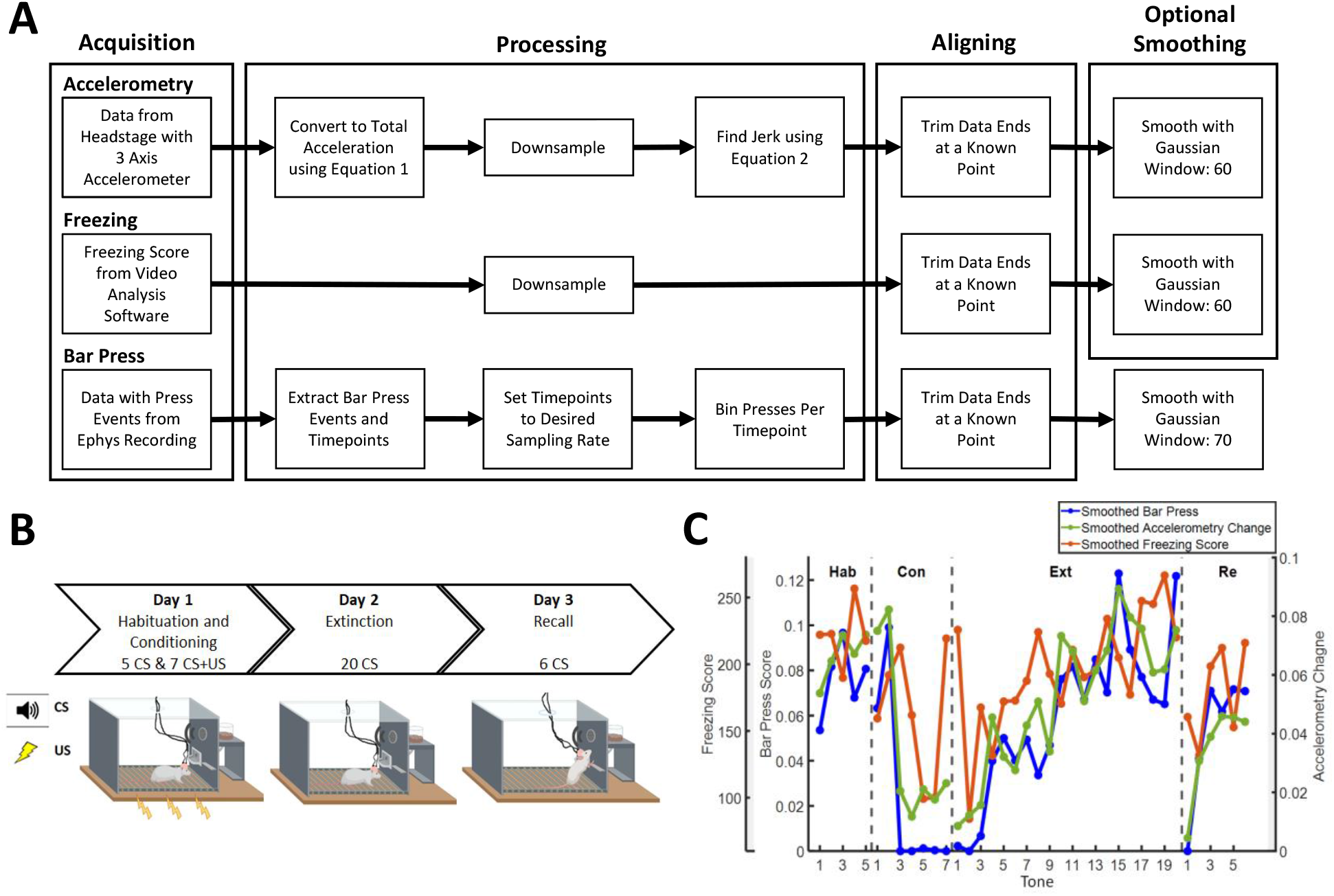
Accelerometry, bar press, and freezing scores follow similar patterns through tone-shock conditioning in rats. A: Signal processing chain used for each data type. B: Three day tone-shock conditioning protocol. Speaker with sound waves represents the conditioned stimulus (CS, tone). Lightning bolts represent the unconditioned stimulus (US, foot shock). Image includes a rat with a headcap, cables, and lever for bar pressing. C: Single animal example of defensive behavior. Each point represents the mean smoothed bar press rate, accelerometry change, or video-derived freezing score during a single tone. All three curves show the same qualitative change in behavior.

Defensive behavior (freezing) was quantified by offline video analysis, to serve as ground truth for an accelerometry-based algorithm. As above, video was captured during all behavioral experiments using a Logitech HD Pro Webcam C910 and attached Neewer Digital high definition .45x super wide-angle lens. Video was recorded using Debut Professional (NCH Software, Canberra, Australia) at 30 frames per second. To link the ground truth to protocols used across many labs, freezing behavior was extracted from the videos via ANY-maze (Stoelting Co.) using the built-in freezing detection feature. A focus region outlining the apparatus was drawn to match the chamber for each video. The software produced a ‘freezing score’ for each frame, based on the change in pixels within the focus region between successive frames. Increased change between frames produced a higher freezing score.

Reward seeking behavior (bar press) was quantified post session as an active measure of defensive behavior, with a decrease in pressing interpreted as increased threat response. Bar press events were logged in the electrophysiology event data via a data acquisition system (DAQ) (USB 6343-BNC or PCIe-6353, National Instruments, Woburn, MA, USA). Presses and timepoints were extracted from the event data for further processing.

### Accelerometry Analysis

During all behavior sessions, accelerometry data were acquired continuously at 30 kHz, using a RHD 2132 electrophysiology head stage with built-in 3-axis accelerometer (Intan Technologies LLC, Los Angeles, CA, USA). Data was recorded through an Open Ephys acquisition system (Open Ephys, Cambridge, MA, USA), a widely used open-source platform for in vivo electrophysiology (Siegle et al., 2014). Tones/trials were marked in the accelerometry data by digital event signals generated from the DAQ. Accelerometry recordings were synchronized to the video by aligning the first ‘tone on’ event in both the accelerometry data and video.

### Data Processing

#### Accelerometry

Data processing used Matlab (Mathworks, Natick, MA). The headstage orientation during recordings was unknown and included sporadic rodent movement. Therefore, total acceleration included both the rodent’s movement and a constant component due to gravity. The X, Y, and Z accelerometer voltages were converted to total acceleration using the following equation:

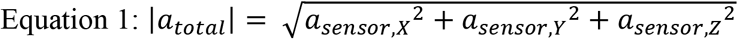

Total acceleration was resampled from the original sampling rate of 30,000 samples/second to 10 samples/second. This and all other resampling used the Matlab resample() function, which includes an anti-aliasing filter. To further identify periods of defensive immobility, we calculated change in total acceleration (jerk) using equation 2, where n indexes a single time point. We compared both *a*_*total*_and *a*_*change*_ to other defensive behavior metrics, as described below.

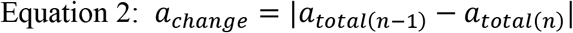

‘Tone on’ and ‘tone off’ events were identified in the electrophysiology ‘events’ file. Both *a*_*total*_and *a*_*change*_ were trimmed 30 seconds before the first ‘tone on’ and after the last ‘tone off’ events to align to the other defensive metric data sets. Total acceleration and acceleration change were then smoothed using a Gaussian filter with a window of 60 samples, which eliminated non-biological noise transients. The filter window width was chosen manually through inspection of the smoothed traces. We compared *a*_*total*_and *a*_*change*_, with and without this smoothing, to other defensive behavior metrics as described below.

#### Video

We ran each video through AnyMaze software, which provided a freezing score and timepoint per frame. Freezing scores and timepoints were resampled at 10 samples/second and trimmed 30 seconds before the first and after the last tone to align with other data. Video was started at the same time as the tone-shock conditioning paradigm, hence tones occurred at a known time point for each recording. Freezing scores were then smoothed with a Gaussian filter with a window of 60 samples. The window width was again chosen manually.

#### Bar Press

Bar press events and continuous time were extracted from the electrophysiology recording. Continuous time was downsampled to 10 samples/second. The number of presses was then binned into each time interval. As they were already aligned, the bar press data were trimmed at the same time points as the total acceleration data. Once trimmed, presses were smoothed with a Gaussian filter of 70 which changed the data from a point process count to an estimation of a continuous press rate/press likelihood. The filter width was optimized visually; a filter width of 70 provided better smoothing than 60 without over-smoothing the bar press response.

#### Behavioral Correlations

Behavior as measured via accelerometry, video based freezing scores, and bar press were assessed for each animal (n = 28). No data was excluded from analysis. Pearson correlations between accelerometry data, video derived freezing scores, and bar press scores were calculated using Matlab. We correlated freezing and bar press against accelerometry as transformed through six processing methods: raw (*a*_*total*_, R), raw change (*a*_*change*_, C), smoothed raw (S), smoothed change (SC), smoothed shifted (SSh), and smoothed change shifted (SCSh). Shifted methods involve shifting the bar press or freezing data by a lag. We determined the lag by computing cross correlation between bar press, freezing, and acceleration, at lags from +20 to -20 samples.

We then set the lag, for each animal and each signal, at the value that maximized cross correlation. The shifted analyses compensated for any potential lag between the three sensors.

## Statistical Analysis

For each animal and for each accelerometry processing option, we calculated the Pearson correlations between metrics as above. We computed a single value for each correlation across the three testing days (Habituation/Conditioning, Extinction, and Recall). To identify whether different accelerometry processing methods differed in their correlations with standard defensive metrics, we performed a repeated measures ANOVA on these correlation values. We considered this as repeated measures because the different processing methods contained overlapping elements, and thus were likely correlated with each other. We compared differences between methods with post hoc Tukey’s HSD testing. To test the overall correlation between our three defensive metrics, we compiled the R values from the best method identified in the ANOVA. For each metric-to-metric comparison (bar press x freezing, freezing x accelerometry, bar press x accelerometry), we used a one-sample t-test to determine whether the mean correlation exceeded zero. All tests were considered statistically significant at p<0.05.

### Results

#### Accelerometry Validation

We applied similar processing chains to our three defensive behavior metrics (Fig. 1A). As an initial validation, we examined defensive behavior as averaged within each 30-second tone during a conditioning and extinction experiment (Fig. 1B). All three metrics showed the same qualitative pattern in individual animals (Fig. 1C): suppression during the Conditioning phase, slow recovery during Extinction (as the animals learned that CS no longer predicted US), and a more rapid recovery during the subsequent Recall test. This is the expected pattern for this behavioral assay (Colom-Lapetina et al., 2019; Gruene et al., 2015; Li et al., 2014; Mueller et al., 2009).

#### Behavioral Correlations

Although Figure 1 suggests that change in accelerometry, bar press rate, and freezing scores measure similar behavioral processes, we found little correlation between these defensive measurements at the more granular, single-timepoint level. Within an individual CS tone presentation, the three measures did not track well (Fig. 3A). Accelerometry and bar press were most correlated (r = 0.53, t(27) = 23.29,p = 0). Freezing and bar press (r = 0.10, t(27) = 9.04, p = 1.19e-9), and freezing and accelerometry were less correlated (r = 0.11, t(27) = 7.91, p = 1.68e-8, Fig. 3B).

There were significant differences between accelerometry processing methods for both freezing (F(5)=5.26,p=1.95e-4) and bar press (F(5)=39.02,p=1.31e-6) correlations. We found no rat and processing method interactions for both freezing (F(5)=0.21,p=0.96) and bar press (F(5)=1.26,p=0.27). Additional processing steps significantly increased correlations between accelerometry and both freezing and bar press (Table 1). The shifting operator improved freezing correlations but had no effect on bar press. The optimal lags from freezing and accelerometry cross correlations (Figure 2), were inconsistent across animal and session. Further, the majority of optimal lags were either 0 (no shifting required) or were at or beyond +/-20 samples. The latter would imply an over 2 second lag which would not be physically reasonable. Therefore, unshifted correlations were used for analyses.

**Table 1:** Tukey HSD p-values for post hoc comparisons. A: Accelerometry processing methods and freezing. B: Accelerometry processing methods and bar press. Abbreviations are defined in Fig. 4.

| Acceleration Method x Freezing |  |  |  | Acceleration Method x Bar Press |  |  |  |
| --- | --- | --- | --- | --- | --- | --- | --- |
| Method 1 | Method 2 | Difference | P-value | Method 1 | Method 2 | Difference | P-value |
| R | S | -0.006 | 1.71e-4** | R | S | -0.026 | 4.68e-4** |
| R | SC | -0.026 | 0.004** | R | SC | -0.251 | 2.07e-8** |
| R | SSh | -0.006 | 1.31e-4** | R | SCSh | -0.231 | 2.09e-8** |
| R | SCSh | -0.027 | 0.002** | RC | SC | -0.234 | 2.07e-8** |
| RC | SC | -0.024 | 0.002** | RC | SCSh | -0.215 | 2.07e-8** |
| RC | SCSh | -0.026 | 8.43e-4** | S | SC | -0.224 | 2.14e-8** |
| S | SC | -0.020 | 0.034* | S | SSh | 0.021 | 0.001** |
| S | SCSh | -0.021 | 0.019* | S | SCSh | -0.205 | 2.53e-8** |
| SC | SSh | 0.020 | 0.027* | SC | SSh | 0.245 | 2.07e-8** |
| SC | SCSh | -0.002 | 2.21e-4** | SC | SCSh | 0.019 | 0.029* |
| SSh | SCSh | -0.022 | 0.015* | SSh | SCSh | -0.226 | 2.08e-8** |

**Figure 2:**
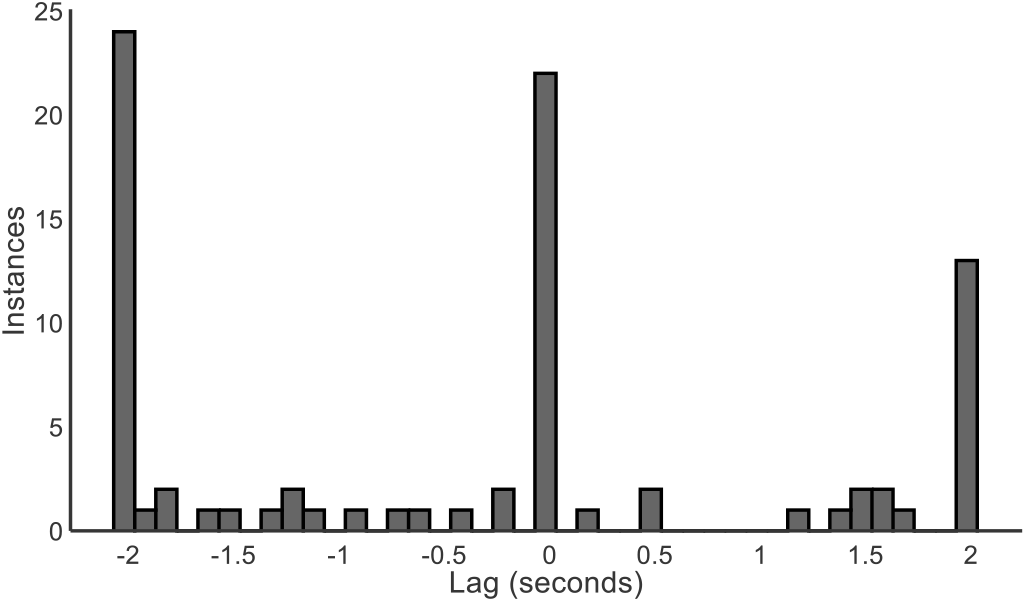
Lags between video based freezing and accelerometry scores are highly inconsistent between animals. The histogram shows, across all rats and sessions (n = 84), the lags that maximized the cross correlation of smoothed freezing and smoothed accelerometry change. The strongest correlations occur either at a lag of 0 (no shift) or at lags beyond 2 seconds that are biologically implausible.

**Figure 3:**
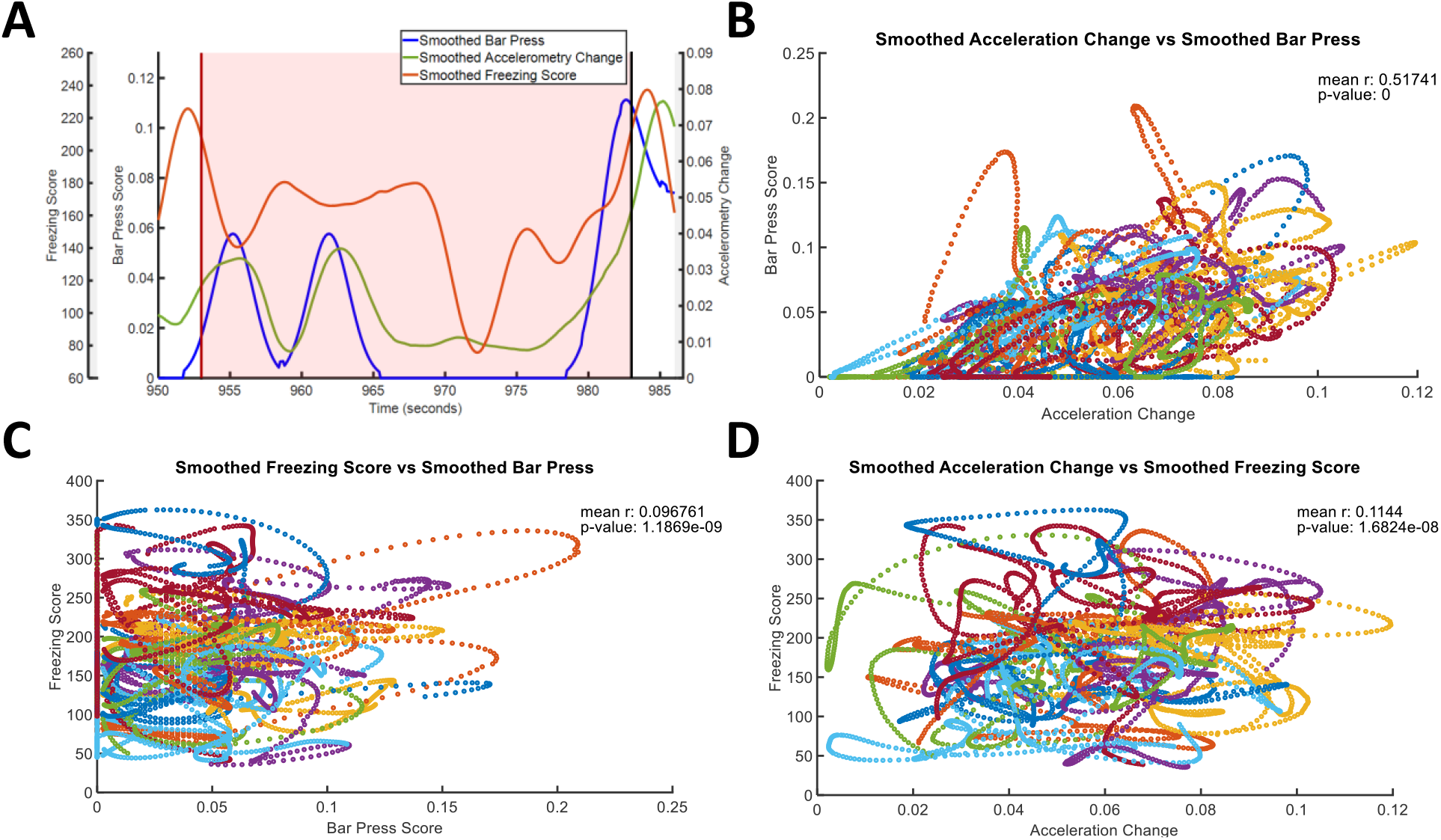
Little correlation between defensive behavior metrics. A: Single animal example of smoothed bar press, accelerometry change, and freezing score during one tone of the Recall phase. Shaded area represents tone duration, with red and black vertical lines representing ‘tone on’ and ‘tone off’ respectively. All three measures suppress during the tone, but in different amounts at different times. B: Scatterplot with mean correlation and p-value of defensive behavior as measured by smoothed bar press and smoothed acceleration change over 200 samples. Each color represents an individual animal for figures B,C, and D. C: Scatterplot with mean correlation and p-value of defensive behavior as measured by smoothed bar press and smoothed freezing score over 200 samples. D: Scatterplot with mean correlation and p-value of defensive behavior as measured by smoothed freezing score and smoothed acceleration change over 200 samples.

**Figure 4:**
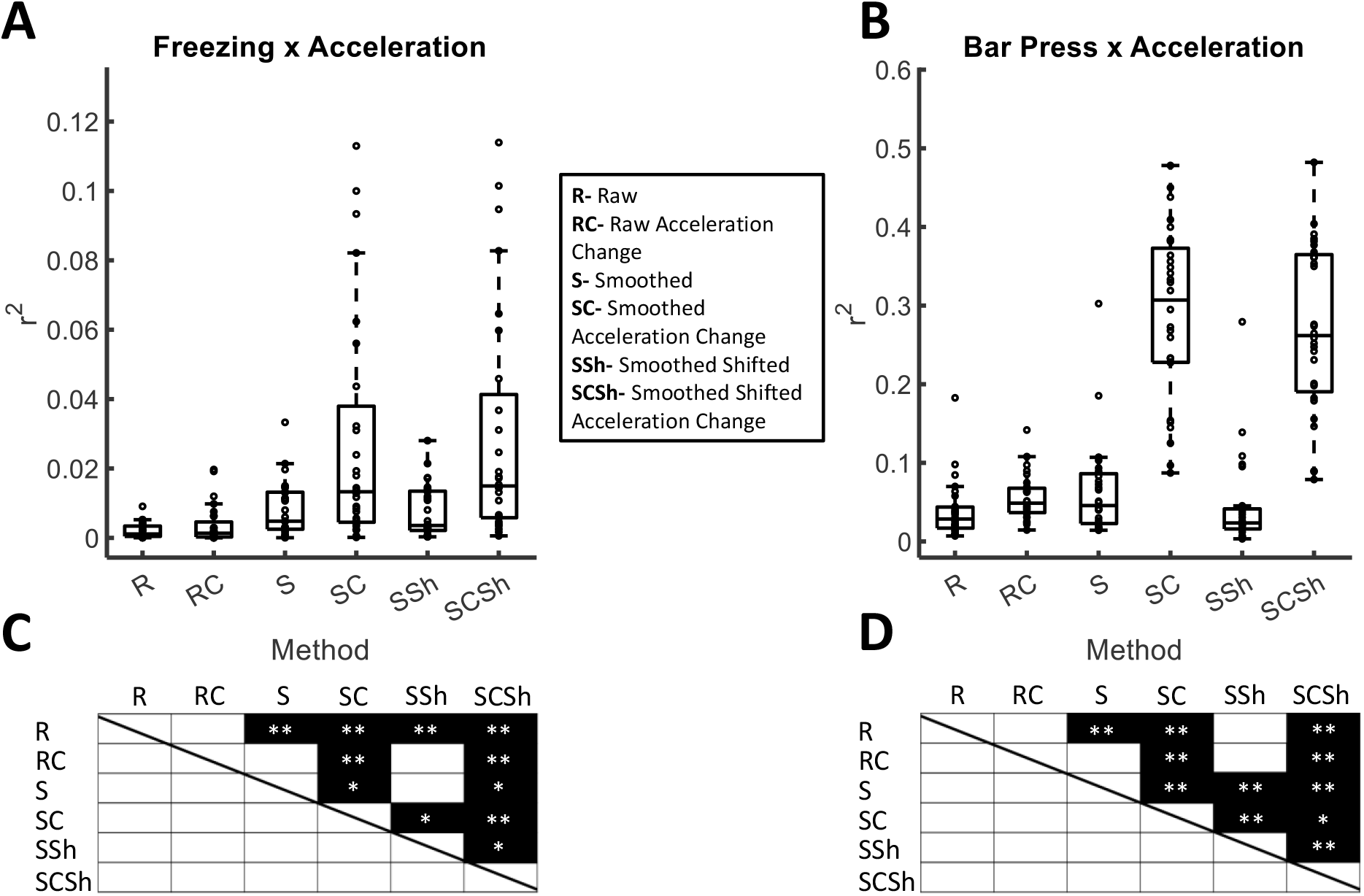
Additional processing increases correlations between accelerometry data and other metrics. A: Correlations between accelerometry processing methods and video derived freezing scores. Freezing scores are raw in R and RC methods and smoothed in all smoothed correlations. Each point within a box is an animal. B: Correlations between accelerometry processing methods and bar press. All bar press scores are smoothed. C, D: Post hoc testing outputs. Each black square represents a significant post-hoc Tukey test. *significant at p<0.05, **significant at p<0.01. C and D correspond to testing using freezing and bar press as the correlated variable, respectively.

## Discussion

In this work, we developed an approach to detect rodent defensive behaviors based on continuous motion sensors. The change in sensor voltage from a built-in 3 axis accelerometer can track behavior, at a coarse trial-to-trial level, comparably to video or bar press suppression methods. As shown in Figure 1C, change in accelerometry follows similar patterns as video derived freezing score and bar press rate. Accelerometry can thus substitute for these metrics in conditioning and extinction paradigms, and in other paradigms where defensive immobility is of interest (e.g., in open fields, elevated mazes, or predator threat paradigms). Accelerometer measurements of defensive behavior require less specialized software and no additional animal training. They may thus be useful in situations where resources are limited.

Further, we performed these measurements through an open source physiology system (Siegle et al., 2015) that supports many plugins for real time signal processing (Schatza et al., 2021; Wodeyar et al., 2021). Thus, events could be triggered based on changes in acceleration, e.g. to initiate or terminate bouts of defensive behavior in a closed loop fashion. Real time accelerometry readings could be combined with other developed algorithms (Pasquet et al., 2016; Venkatraman et al., 2010) to assess other, non-defensive behaviors such as grooming or reaction time. In this regard, accelerometry may have advantages over real-time video tracking, as it requires few data channels at a lower sampling rate, enabling faster closed loop processing and operation in more resource-limited environments. Similarly, accelerometry may be valuable in environments where animals are not constrained to a small operant arena, but can freely explore complex environments (e.g., using wireless recording (Yin & Hanson, n.d.). This has led to proposals to use accelerometry alone as, e.g., a readout of pathologic human behavior (Zamora et al., n.d.).

An important caveat is that the modest correlations between all three metrics at the finer timescale suggest that they capture distinct elements of defensive behavior. Time lag cannot account for the modest correlations between accelerometry and bar press. Although shifting the freezing time series did significantly increase its correlation with accelerometry, the observed lags were inconsistent and more conducive to noise than predictive lag between metrics. Accelerometry might have value as an independent metric of defensive behavior, but does not seem to reflect aspects of defensive processes that occur before visible freezing.

This initial proof of concept does have limitations. First, most studies of defensive behavior using video tracking define thresholds, below which an animal is considered freezing. We did not develop these thresholds in this work, but future research could attempt to dichotomize behaviors. Second, we focused on comparisons to freezing, but some rodents also display rapid escape behaviors (“darting”) in response to threat (Colom-Lapetina et al., 2019; Gruene et al., 2015). Accelerometry may be more reliable than video at detecting such behaviors, but the relevant processing algorithms will need to be developed. Third, we did not account for the unknown gravity vector in the accelerometer calculations. More accurate movement and tilt estimations can be made by including this vector. For studies looking into more complex behavior, either including the gravity vector or using a recording device that includes a gyroscope (*Digital Headstage Cereplex µ*, n.d.; *Next Gen. Acquisition System*, n.d.; Fayat et al., 2021) may be warranted.

Despite these limitations, this work shows that accelerometry is a viable method of measuring defensive behavior and carries information not found in other defensive assays. It thus may become an important part of a behavioral neuroscience toolkit.

## Code Availability

At the time of publication, both an analysis script to perform accelerometry smoothing/timeseries extraction and code to reproduce the figures in this manuscript will be available at https://github.com/tne-lab.

## Acknowledgements

This work was supported by grants R21MH109722, R21MH113103, and R01MH11938402 from the National Institutes of Health, by the MnDRIVE Brain Conditions Initiative, and the Minnesota Medical Discovery Team on Addictions.

## References

Amorapanth, P., Nader, K., & Ledoux, J. E. (1999). Lesions of Periaqueductal Gray Dissociate-Conditioned Freezing From Conditioned Suppression Behavior in Rats. Learning & Memory, 6(5), 491. https://doi.org/10.1101/LM.6.5.491

Anagnostaras, S. G., Wood, S. C., Shuman, T., Cai, D. J., LeDuc, A. D., Zurn, K. R., Zurn, J. B., Sage, J. R., & Herrera, G. M. (2010). Automated assessment of Pavlovian conditioned freezing and shock reactivity in mice using the VideoFreeze system. Frontiers in Behavioral Neuroscience, 4(SEP). https://doi.org/10.3389/fnbeh.2010.00158

Campos, A. C., Fogaça, M. V., Aguiar, D. C., & Guimarães, F. S. (2013). Animal models of anxiety disorders and stress. Revista Brasileira de Psiquiatria, 35(SUPPL.2). https://doi.org/10.1590/1516-4446-2013-1139

Colom-Lapetina, J., Li, A. J., Pelegrina-Perez, T. C., & Shansky, R. M. (2019). Behavioral diversity across classic rodent models is sex-dependent. Frontiers in Behavioral Neuroscience, 13. https://doi.org/10.3389/fnbeh.2019.00045

Deslauriers, J., Toth, M., Der-Avakian, A., & Risbrough, V. B. (2018). Current Status of Animal Models of Posttraumatic Stress Disorder: Behavioral and Biological Phenotypes, and Future Challenges in Improving Translation. Biological Psychiatry, 83(10), 895–907. https://doi.org/10.1016/j.biopsych.2017.11.019

Digital Headstage Cereplex µ. (n.d.). DigitalOne. Retrieved December 1, 2021, from https://www.brainlatam.com/manufacturers/headstages-ephys/digital-headstage-cereplex-%C2%B5--338

Fayat, R., Betancourt, V. D., Goyallon, T., Petremann, M., Liaudet, P., Descossy, V., Reveret, L., & Dugué, G. P. (2021). Inertial measurement of head tilt in rodents: Principles and applications to vestibular research. Sensors, 21(18). https://doi.org/10.3390/s21186318

Gruene, T. M., Flick, K., Stefano, A., Shea, S. D., & Shansky, R. M. (2015). Sexually divergent expression of active and passive conditioned fear responses in rats. ELife, 4. https://doi.org/10.7554/elife.11352

Li, Y., Dong, X., Li, S., & Kirouac, G. J. (2014). Lesions of the posterior paraventricular nucleus of the thalamus attenuate fear expression. Frontiers in Behavioral Neuroscience, 8(MAR). https://doi.org/10.3389/FNBEH.2014.00094

Liu, X. L., Yu, S. Y., Flierman, N. A., Loyola, S., Kamermans, M., Hoogland, T. M., & Zeeuw, C. I. D. (2021). OptiFlex: Multi-Frame Animal Pose Estimation Combining Deep Learning With Optical Flow. Frontiers in Cellular Neuroscience, 15. https://doi.org/10.3389/fncel.2021.621252

Lo, M. C., Younk, R., & Widge, A. S. (2020). Paired Electrical Pulse Trains for Controlling Connectivity in Emotion-Related Brain Circuitry. IEEE Transactions on Neural Systems and Rehabilitation Engineering, 28(12). https://doi.org/10.1109/TNSRE.2020.3030714

Mathis, A., Schneider, S., Lauer, J., & Mathis, M. W. (2020). A Primer on Motion Capture with Deep Learning: Principles, Pitfalls, and Perspectives. Neuron, 108(1), 44–65. https://doi.org/10.1016/j.neuron.2020.09.017

Mueller, D., Olivera-Figueroa, L. A., Pine, D. S., & Quirk, G. J. (2009). The effects of yohimbine and amphetamine on fear expression and extinction in rats. Psychopharmacology, 204(4), 599. https://doi.org/10.1007/S00213-009-1491-X

Next gen. acquisition system: ONIX. (n.d.). Open Neuro Interface. Retrieved December 1, 2021, from https://open-ephys.org/next-gen-acquisition-system

Pasquet, M. O., Tihy, M., Gourgeon, A., Pompili, M. N., Godsil, B. P., Léna, C., & Dugué, G. P. (2016). Wireless inertial measurement of head kinematics in freely-moving rats. Scientific Reports, 6. https://doi.org/10.1038/srep35689

Robinson, O. J., Pike, A. C., Cornwell, B., & Grillon, C. (2019). The translational neural circuitry of anxiety. Journal of Neurology, Neurosurgery & Psychiatry, 90(12), 1353–1360. https://doi.org/10.1136/jnnp-2019-321400

Schatza, M. J., Blackwood, E. B., Nagrale, S. S., & Widge, A. S. (2021). Toolkit for Oscillatory Real-time Tracking and Estimation (TORTE). Journal of Neuroscience Methods, 109409. https://doi.org/10.1016/J.JNEUMETH.2021.109409

Siegle, J. H., Hale, G. J., Newman, J. P., & Voigts, J. (2015). Neural ensemble communities: Open-source approaches to hardware for large-scale electrophysiology. Current Opinion in Neurobiology, 32, 53–59. https://doi.org/10.1016/j.conb.2014.11.004

Sousa, N., Almeida, O. F. X., & Wotjak, C. T. (2006). A hitchhiker’s guide to behavioral analysis in laboratory rodents. Genes, Brain and Behavior, 5(2), 5–24. https://doi.org/10.1111/j.1601-183X.2006.00228.x

Terburg, D., Scheggia, D., Rio, R. T. del, Klumpers, F., Ciobanu, A. C., Morgan, B., Montoya, E. R., Bos, P. A., Giobellina, G., Burg, E. H. van den, Gelder, B. de, Stein, D. J., Stoop, R., & Honk, J. van. (2018). The Basolateral Amygdala Is Essential for Rapid Escape: A Human and Rodent Study. Cell, 175(3). https://doi.org/10.1016/j.cell.2018.09.028

Venkatraman, S., Jin, X., Costa, R. M., & Carmena, J. M. (2010). Investigating neural correlates of behavior in freely behaving rodents using inertial sensors. Journal of Neurophysiology, 104(1). https://doi.org/10.1152/jn.00121.2010

Venkatraman, S., Long, J. D., Pister, K. S. J., & Carmena, J. M. (2007). Wireless inertial sensors for monitoring animal behavior. Annual International Conference of the IEEE Engineering in Medicine and Biology - Proceedings. https://doi.org/10.1109/IEMBS.2007.4352303

von Ziegler, L., Sturman, O., & Bohacek, J. (2021). Big behavior: Challenges and opportunities in a new era of deep behavior profiling. Neuropsychopharmacology, 46(1), 33–44. https://doi.org/10.1038/s41386-020-0751-7

Wodeyar, A., Schatza, M., Widge, A. S., Eden, U. T., & Kramer, M. A. (2021). A state space modeling approach to real-time phase estimation. ELife, 10. https://doi.org/10.7554/eLife.68803

Yin, A., & Hanson, T. (n.d.). GitHub—Allenyin/allen_wireless: Wireless recording system, forked from Tim Hanson’s myopen project. [PostScript, Assemely, C, C++, Logos, Python]. https://github.com/allenyin/allen_wireless

Zamora, M., Toth, R., Morgante, F., Ottaway, J., Gillbe, T., Martin, S., Lamb, G., Noone, T., Benjaber, M., Nairac, Z., Constandinou, T. G., Herron, J., Aziz, T. Z., Gillbe, I., Green, A. L., Pereira, E. A. C., & Denison, T. (n.d.). DyNeuMo Mk-1: Design and Pilot Validation of an Investigational Motion-Adaptive Neurostimulator with Integrated Chronotherapy. https://doi.org/10.1101/2020.09.10.292284

